# Liver transcriptomic and methylomic analyses identify transcriptional MAPK regulation in facultative hibernation of Syrian hamster

**DOI:** 10.1101/2022.12.01.518631

**Authors:** Marloes M. Oosterhof, Louis Coussement, Victor Guryev, Vera A. Reitsema, Jojanneke J. Bruintjes, Maaike Goris, Hjalmar R. Bouma, Tim de Meyer, Marianne G. Rots, Robert H. Henning

## Abstract

Hibernation consist of alternating torpor/arousal phases, during which animals cope with repetitive hypothermia and ischemia-reperfusion. Due to limited transcriptomic and methylomic information for facultative hibernators, we here conducted RNA and whole genome bisulfite sequencing in liver of hibernating Syrian hamster *(Mesocricetus auratus*). Gene Ontology analysis was performed on 844 differentially expressed genes (DEGs) and confirmed the shift in metabolic fuel utilization, inhibition of RNA transcription and cell cycle regulation as found in seasonal hibernators. We show a so far unreported suppression of MAPK and PP1 pathways. Notably, hibernating hamsters showed upregulation of MAPK inhibitors (DUSPs and SPRYs) and reduced levels of MAPK induced transcription factors. Promoter methylation was found to modulate the expression of genes targeted by these transcription factors. In conclusion, we document gene regulation between hibernation phases, which may aid the identification of pathways and targets to prevent organ damage in transplantation or ischemia-reperfusion.

## Introduction

Hibernation is an adaptive strategy to cope with inadequate energy supply because of low food availability or challenging thermoregulatory conditions and is characterized by metabolic suppression, lowering of body temperature (T_*b*_) and cessation of locomotive activity during periods of torpor. Torpor periods are alternated with briefer arousal periods, an energetically expensive process restoring metabolism and T_*b*_. Whereas hibernators tolerate the repetitive, drastic alterations in physiology during torpor-arousal cycles, similar alterations result into organ dysfunction and damage in non-hibernators such as rat and humans^1,2^.

Numerous studies of hibernating mammals have revealed changes in gene expression. Transcriptome analysis across hibernation phases mainly in seasonal, i.e. non-hoarding squirrels, identified a switch in expression of metabolic genes to accommodate fatty acid oxidation throughout hibernation^3–6^. In contrast to seasonal hibernators, Syrian hamster (*Mesocricetus auratus*) is a facultative hibernator, which enters hibernation in response to environmental cues, such as lowering of ambient temperature and shortening of daylight, rather than being driven by an endogenous circannual rhythm^7^. Importantly, their facultative nature of hibernation might identify particular pathways that may be exploited to prevent organ damage in the human setting. As no comprehensive transcriptomic studies in facultative hibernating species have yet been reported, we examined expression changes, as well as DNA methylation differences, in hibernating Syrian hamster. To investigate the metabolic aspect of facultative hibernation, we studied the liver of the Syrian hamster. Liver is considered a crucial organ in hibernation, as it accommodates the bulk of the metabolic changes from summer to hibernation^8^. Additionally, liver is the site of synthesis of enzymes involved in gluconeogenesis and ketone body formation, processes required for fuel generation during the hibernation season^9^. Therefore, we performed unbiased RNA sequencing and DNA methylation analysis in liver from summer, torpid and arousing Syrian hamster to explore mechanisms of hibernation initiation and organ protection in a facultative hibernator, which shows suppression of the mitogen-activated protein kinase (MAPK) pathway in torpor and upregulated promoter methylation in MAPK transcription factor target genes in arousal.

These results provide additional evidence for the relevance of these TFs, as the limited enrichment of their targets may be explained by the inhibitive effect of DNA methylation for non-responding TF target genes.

## Results

### Analysis of gene expression differences between hibernation stages

Liver gene expression changes during the torpor/arousal cycle were analyzed by comparing RNA-sequencing (RNA-seq) data of summer euthermic (SE) animals and the two hibernation stages, torpor late (TL) and arousal early (AE). The number of differentially expressed genes (DEGs) was 272 for SE vs TL, 137 for TL vs AE and 435 for SE vs AE (Fig.1A, Table 2). Number and overlap of genes during hibernation stages are summarized in a Venn diagram (Fig.1A). Of the DEGs, 394 were unique for a single comparison between stages, 225 were differentially expressed for two comparisons, whereas none of the DEGs were differentially expressed for all three contrasts (Fig.1A).

**Fig. 1.**
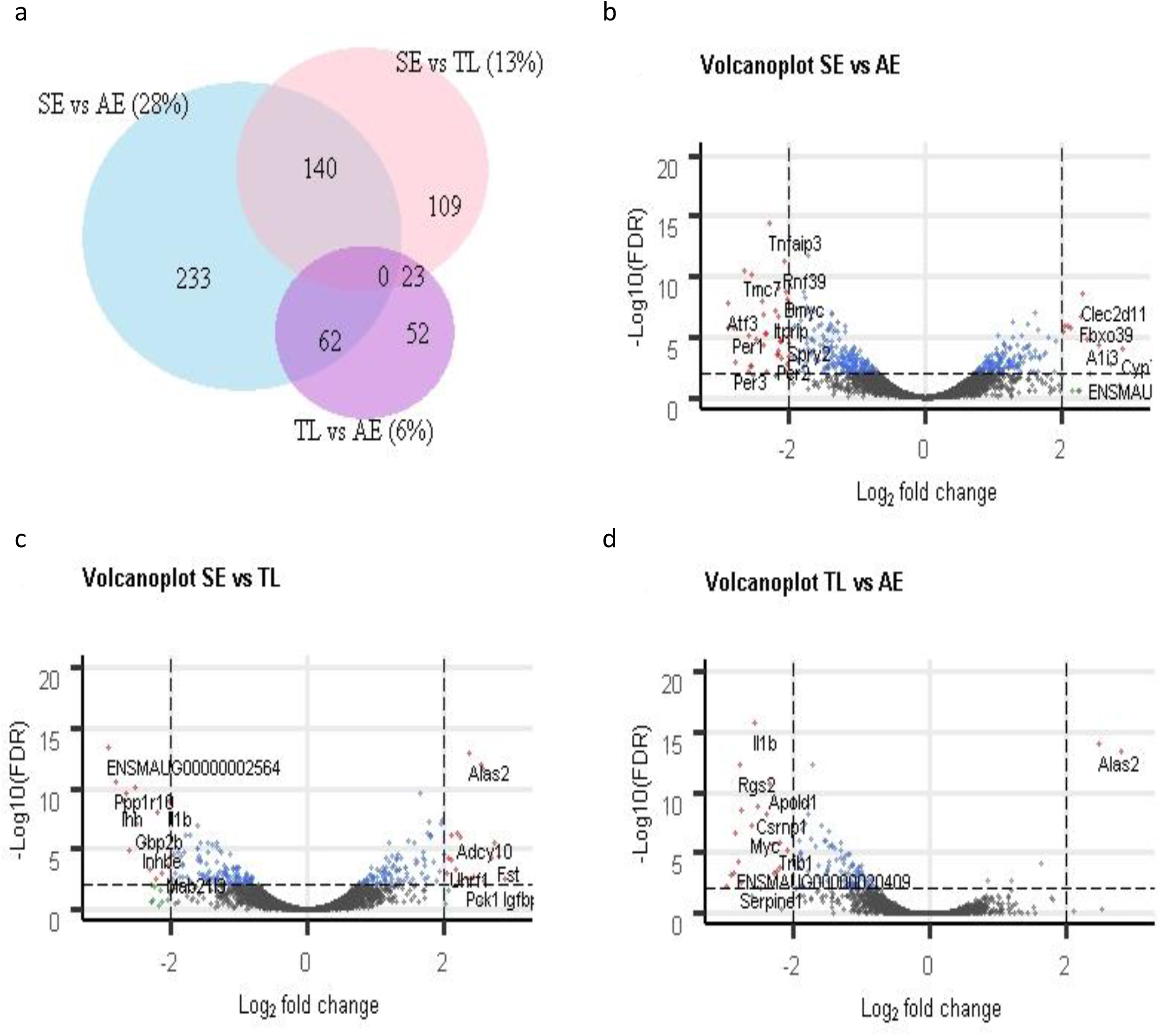
Differentially expressed genes (DEGs) in the RNA-sequencing data. a. Venn diagram of DEGs (FDR < 0.01) identified SE vs TL, TL vs AE and SE vs AE. b-d. Volcano plots of genes identified using RNA-seq for SE vs TL, TL vs AE and SE vs AE respectively. The FDR cut-off line was FDR < 0.01 (vertical dashed line). Vertical dashed lines represent cut-off of logFC of -2 and +2

Next, the distribution of up- and downregulated DEGs was examined. In SE vs TL and SE vs AE, the number of up- and downregulated genes was almost equal, with a small excess of downregulated genes when comparing SE to TL (55%), and of upregulated genes when comparing SE to AE (60%, Table 2). In striking contrast, the transition from TL to AE was characterized almost exclusively by upregulation of gene expression in AE, representing 93% of the 137 DEGs (P < 2.2E-16, binomial test). Of the 128 upregulated DEGs between TL and AE, 63 were also upregulated in AE compared to SE, thus representing genes that are overexpressed specifically upon arousal.

The unique DEGs of SE vs TL, TL vs AE and SE vs AE were visualized by Volcano plots (Fig. 1B-D, Table S1) as well as a heatmap for all three pairwise comparisons (Fig. 2), demonstrating large expression differences between stages. The heatmap identified five different clusters of genes of which cluster 1 (301 DEGs) and cluster 5 (256 DEGs) represent genes being upregulated in SE compared to AE and TL, respectively. In cluster 2, 54 genes are downregulated in TL compared to other groups, while cluster 4 (13 DEGs) represents genes exclusively upregulated in TL. Finally, cluster 3 represents 94 genes that are downregulated in TL compared to SE and AE.

**Fig. 2.**
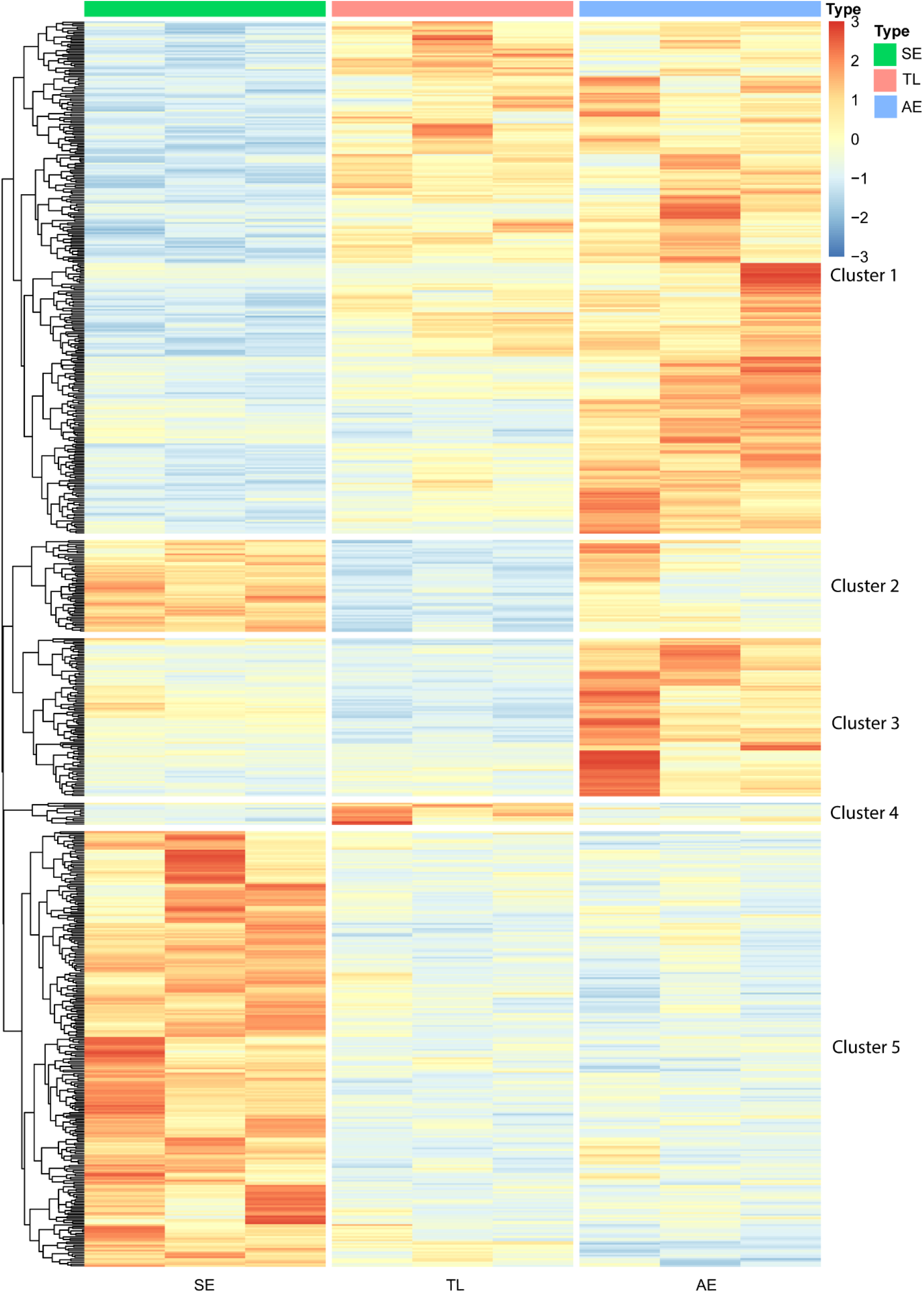
Hierarchical clustering of differentially expressed genes (DEGs; FDR<0.01). Five clusters were formed using Euclidean complete linkage clustering as depicted with 1-5.

Comparing SE to TL, two upregulated genes in TL had a remarkably low FDR and high fold change: pro-platelet basic protein and tubulin beta-1 chain (*PPBP*: 20-fold, *TUBB1*: 12-fold). Both genes belong to cluster 4, which only contains 13 genes. These two genes are mainly expressed in platelets, and their upregulation is in line with the storage of platelets in liver during torpor^10^. Also 5-aminolevulinate synthase (*ALAS2*) is a prominently upregulated gene in torpor compared to SE and AE (5-aminolevulinate synthase: 5-fold). *ALAS2* encodes an enzyme which catalyzes the first step in the heme biosynthetic pathway, although translation is dependent on adequate iron supply^11^. Three genes in the top 10 of most significantly downregulated genes in TL vs SE (ranking #2, 4 and 7 of the downregulated genes) encode regulatory subunits of protein phosphatase 1 (PP1) (*PPP1R3C* and *PPP1R10* (both 6-fold downregulated) and *PPP1R3B* (5-fold downregulated)). These subunits remained downregulated in AE compare to SE. In liver, PP1 accelerates glycogen synthesis and coordinates carbohydrate storage^12^. Downregulation of these subunits in torpor may thus contribute to the shift from glucose to fatty acid metabolism as part of metabolic rewiring in hibernation.

Cluster 5 represents DEGs that were downregulated in both TL and AE compared to SE, largely representing innate immune response genes and three benzaldehyde dehydrogenase [NAD(P)+] activity genes. The immune response is suppressed during torpor through a strong reduction in circulating leukocyte numbers^13^, decreased phagocytic capacity and complement activity^14^, with rapid restoration of these processes during arousal. Thus, our data support the notion that the immune system in liver of Syrian hamster remains suppressed during early arousal and is reactivated only late in arousal or even after hibernation^15^. This view is consistent with increased expression of Tumor necrosis factor, alpha-induced protein 3 (*TNFAIP3)* in AE, a strong inhibitor of the Toll-like receptor (TLR) pathway and protective against cell death in renal cold ischemia/reperfusion injury^16^.

### Gene Ontology of DEGs

To identify cellular pathways associated with DEGs between specific hibernation stages, GO and KEGG pathways analyses were performed (Fig.1A, Table S2).

First, to identify processes regulating initiation and maintenance of hibernation, up- and downregulated genes in SE vs both TL and AE were examined (clusters 1 and 5 of Fig. 2). Expectedly, differentially regulated pathways include metabolic pathways (e.g. lipid and glucose response) and hormonal pathways governing metabolism (e.g. insulin pathway) (Fig. 3A). Many hibernating species shift from glucose to fatty acid metabolism when entering torpor^5^. GO analysis validated this metabolic shift towards lipid combustion in TL compared to SE, as pathways including “response to lipid” and “cellular response to hormone/insulin stimulus” were covered by the DEGs in SE vs TL (Fig. 3B). In addition, KEGG pathway analyses suggested insulin resistance as a well-represented process in SE vs TL animals (Table S2). In homeostatic environments, insulin helps control blood glucose levels, promoting blood glucose uptake and storage as glycogen. During hibernation, however, animals become relatively resistant to insulin^8^, shifting the metabolic processes from glycolysis to glycogenolysis and gluconeogenesis as important sources for glucose in torpor. This is indeed reflected by the upregulation of a variety of genes in TL, including *GAA*, Glycogen phosphorylase (*PYG)*, Peroxisome proliferator-activated receptor gamma coactivator 1-alpha (*PPARGC1*_α_), Phosphoenolpyruvate Carboxykinase 1 (*PCK1)* and Glucose 6-phosphate (*G6P)*. In addition, glycolysis is inhibited during torpor by the downregulation of the gene encoding the rate limiting enzyme, germinal center kinase 1 (*GCK1)*. Upregulation of pyruvate dehydrogenase kinase 4 (*PDK4*) in TL inhibits pyruvate dehydrogenase and limits conversion of pyruvate into acetyl-CoA, thus repressing glucose derived oxidative phosphorylation in mitochondria and supporting fat combustion, as shown previously in fasted and starved mammals^17^.

**Fig. 3.**
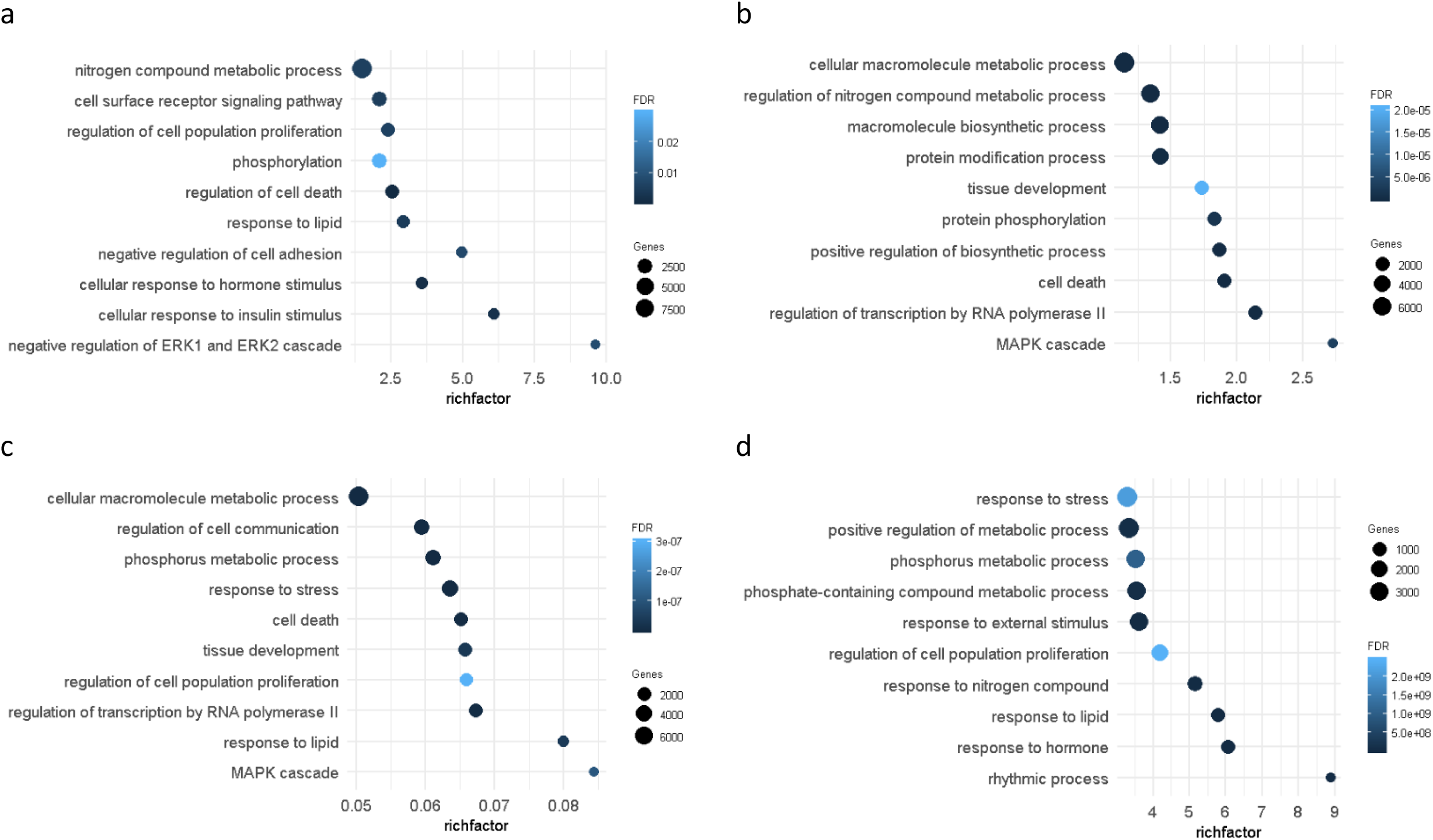
Top 10 GO terms with highest rich factor of differentially expressed genes (DEGs) with FDR < 0.01. # of genes indicate the number of genes found in the GO respective term and the rich factor reflects the proportion of genes in a given pathway. The color indicates the FDR of the term. a. Summer euthermic versus Torpor Late, b. Torpor Late versus Arousal Early, c. Summer euthermic versus Arousal Early, d. Summer euthermic versus Torpor Late and Arousal Early

Although metabolic genes clearly undergo substantial regulation during hibernation, the most striking regulation in all contrasts between stages was observed in the MAPK pathway, particularly its Mitogen-Activated Protein Kinase 1/3 (ERK1/2) cascade (Figs. 2, 4). The DEGs involved represent a major part of cluster 1, which contains upregulated DEGs in both TL and AE versus SE, comprising upregulation of inhibitors of the MAPK pathway (*DUSP1, DUSP4, DUSP5, DUSP8, DUSP10, SPRY1*, and *SPRY2*; Fig. 4). Further, the MAPK pathway is also represented in cluster 3 (e.g. Sprouty Related EVH1 Domain Containing 2 (*SPRED2)*, Ephrin type-A receptor 4 (*EPHA4*)), suggesting differential regulation of specific downstream cascades during hibernation, including the p38 mitogen-activated protein kinase (p38MAPK), extracellular signal-regulated kinase (ERK1/2), and c-Jun-terminal kinase (JNK)^18^. Collectively, GO analysis of gene expression implies a substantial inhibition of the MAPK pathway in TL vs SE (Fig. 4). Consistently, KEGG analysis confirmed regulation of the MAPK pathway when comparing SE vs AE animals (Table S2). The MAPK pathway relays mitogenic signals promoting cell division and proliferation. The suppression of the ERK1/2 cascade members in torpor likely relates to the arrest of the cell cycle, as denoted by upregulation of cyclin-dependent kinase inhibitor 1 (*P21*^*CIP1*^) and of cell death regulating genes Caspase 3 (*CASP3)* and Programmed death-ligand 1 (*PDL1)*. Such proposition is in line with cell cycle arrest in ground squirrel liver by reduced cyclin D and E protein levels and upregulation of cyclin-dependent kinase inhibitors Cyclin-dependent kinase 4 inhibitor B (*P15*^*INK4b*^)and *P21*^*CIP1* 7^.

**Fig. 4.**
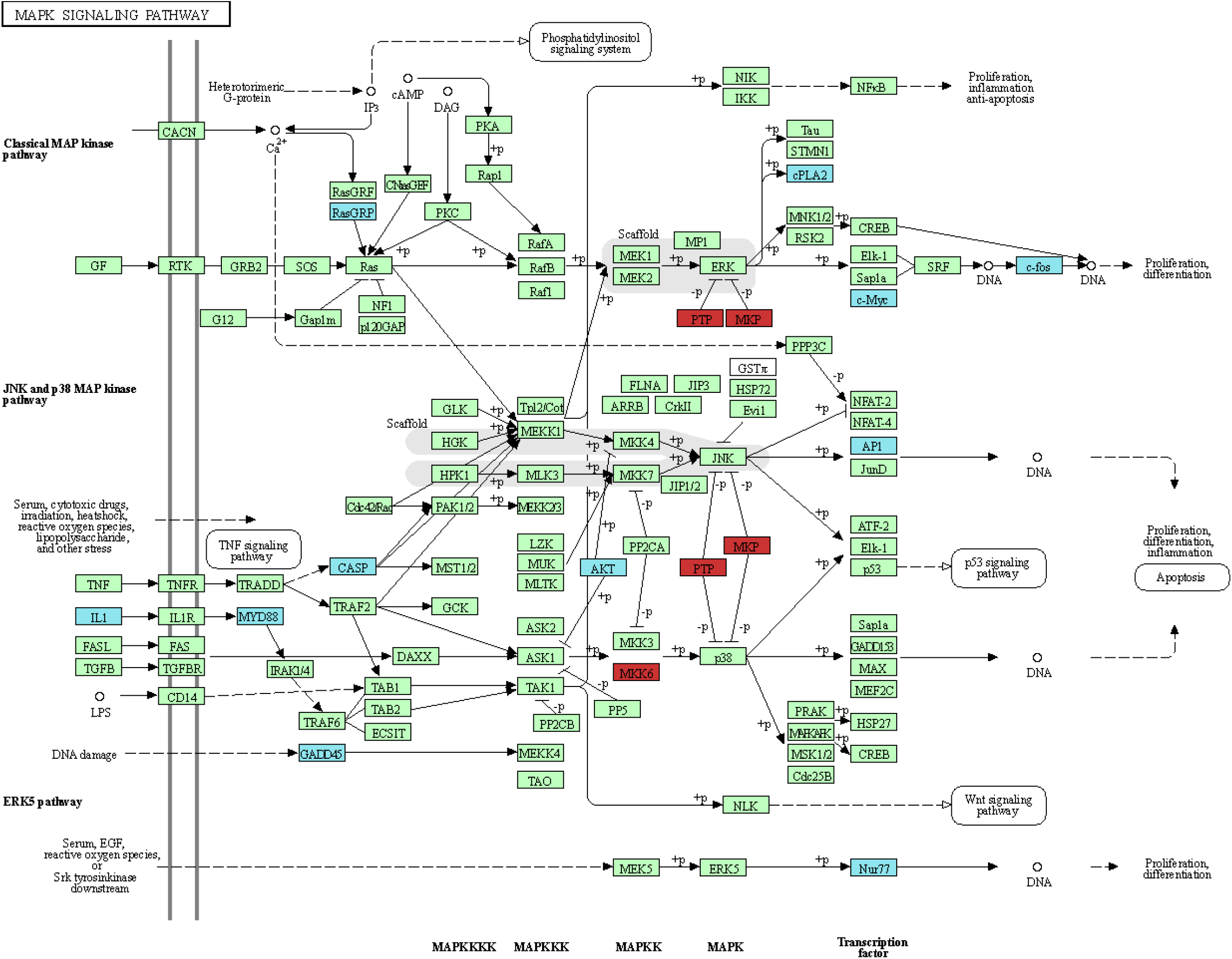
MAPK pathway (KEGG) with differentially expressed genes (DEGs) in torpor (TL) compared to summer euthermic (SE) or arousal early (AE). Red = upregulated in torpor, blue = downregulated in torpor, green = not differentially expressed or not detected.

### GO analysis reveals repression of phosphorylation and RNA expression in torpor

GO analysis uncovered phosphorylation (terms: [protein]phosphorylation and phosphorus metabolic process, Fig. 3A-D) as a recurring pathway in all contrasts. As aforementioned, members of large phosphatase families, such as PP1 subunits, were differentially expressed in torpor. Cluster 2 (Fig. 2, showing downregulated DEGs in TL with upregulation in AE), has a clustering of *PP2A* genes, including Protein Tyrosine Phosphatase Non-Receptor Type 1 (*PTPN1)*, beta-2-adrenergic receptor (*ADRB2)*, and Smg5 nonsense mediated mRNA decay factor (*SMG5)*. PP1 and PP2a are large phosphatase families that are responsible for most of the serine/threonine dephosphorylations, controlling an array of processes, including metabolism and cell cycle. Expression of DEGs related to “regulation of transcription by RNA polymerase II” showed a mildly reduced expression in torpor and strong increase in arousal (cluster 3, Fig. 2). RNA polymerase II is a multiprotein complex and one of the three eukaryotic nuclear RNA polymerases that transcribes DNA into precursors of messenger RNA (mRNA), most small nuclear RNA (snRNA) and microRNA. GO:CC reflects activation of the gene transcription machinery by terms including “nucleus”, “nuclear part” and “nuclear lumen”, consistent with the previously reported reduction of RNA transcription in torpor^19^. Collectively, our data indicated that RNA transcription in hamster is reduced during torpor, followed by an overexpression of RNA transcription in AE.

### GO terms specific for one hibernation stage

DEGs in AE were enriched in “Regulation of cell population proliferation” and “Cell death” when compared with SE. Cell death includes apoptosis, necrosis and autophagy. Of note, a large fraction of DEGs within these terms is related to (extrinsic) apoptotic pathways, suggesting this is a key mechanism activating the apoptosis pathway in arousing hamster liver. In hibernating hamster, an increase in cell damage marker abundance was found in torpid lung^20^, which was rapidly reversed upon arousal. The latter is also suggested by observations in seasonal hibernators, such as the ground squirrel, who arrest cell proliferation in torpor without increased levels of cell death ^7^. Lastly, in SE vs AE animals, “Metabolic responses” and “Nitrogen metabolic compound” are among the most enriched gene ontology terms. These show increased activity during early arousal, possibly making up for protein damage or loss during the torpor stage. Nitrogen metabolic processes generally reflect protein degradation. In ground squirrels, it has been shown that ubiquitination of proteins continues in torpor, whereas proteolysis is inhibited, which may result in increased protein degradation in arousal^21^. Likewise, in Syrian hamster, autophagy is enhanced during early arousal in heart tissue^22^, presumably to clear damaged proteins.

### Differentially expressed genes show minimal promoter DNA methylation differences

To interrogate whether differential DNA methylation represents an underlying factor driving expression changes throughout hibernation, we performed whole genome bisulfite sequencing on liver DNA of these Syrian hamsters. Differentially methylated region (DMR) analysis of the differentially expressed genes (2.5 kb promoter and within-gene regions; +/- 4.5% of all loci) resulted in respectively 49, 57 and 45 clusters (containing ≥ 2 CpGs) retained for the contrasts SE vs TL, SE vs AE and TL vs AE (Table S5). After additional filtering on DMRs overlapping with promoter regions and featuring an average methylation difference of >20%, only (i) 3 DMRs were retained for SE vs TL: Homeodomain Interacting Protein Kinase 2 (*HIPK2)*, Secreted and transmembrane protein 1A (*SECTM1a)* and Aquaporin 3 (*AQP3*); (ii) none for SE vs AE and (iii) 4 DMRs for TL vs AE: *GCK*, Smad Nuclear Interacting Protein 1 (*SNIP1)* and *GM14137* (and one gene without gene symbol: ENSMAUG00000016923). These results demonstrate at most limited evidence for DNA methylation of promotor regions as driving force of differential gene expression.

### Transcription factor (TF) binding site analysis identifies candidate regulatory TFs in arousal animals

Next, to evaluate whether expression of the identified DEGs is driven by key transcription factors, we performed transcription factor binding site (TFBS) analysis on the 2.5 kb promoter regions of differentially expressed genes. As TFBS are well-conserved, the extensive dataset of human TFBS was used, which may also point towards the human homologue (see Material and Methods). TFBS analyses identified three motifs enriched being Early Growth Response 1 (EGR1)-like, MAX Network Transcriptional Repressor (MNT)-like and MYC-like TFs. Interestingly, these TFBS were found to be over-represented in promoters of genes which were upregulated in AE. Also the MAPK target CAMP Responsive Element Binding Protein 1 (CREB1) was found amongst the top enriched TFBS in promoter regions of genes upregulated in arousal animals compared to euthermic animals (52% enrichment). Though not found to be differentially expressed itself, CREB1 has been described as an important target of the p38 MAPK pathway in hibernating bats, suggesting its activation through posttranslational phosphorylation^23^.

Interestingly, EGR1 RNA expression was upregulated in AE compared to TL and SE animals (resp. 5.6 and 18.5 fold, FDR = 9.0E-3 and 4.5E-7). The enrichment of EGR1 TFBS in the promoter regions of genes upregulated in arousal animals compared to both torpor and summer euthermic animals was 39% and 19%, respectively. Also for EGR3 and EGR4 (TFs with highly similar probability weight matrices (PWMs, i.e. binding patterns) as EGF1), higher enrichment of their binding sequences was found (see Fig. S2). Differential expression of EGR3 and EGR4 was not assessed as these genes were filtered out due to too low coverage. Secondly, RNA expression of both MNT and MYC (discussed together due to similar PWM) showed overexpression in arousal animals versus euthermic animals (resp. 1.8 and 12.6 fold, FDR = 1.2E-3 resp. 4.9E-14) and their TFBS enrichment was significant (resp. 41% and 25%) in promoter regions of genes overexpressed in arousal animals compared to euthermic animals. Additionally, two related TFs were found enriched with highly similar PWMs: MYCN and MAX (see Fig. S2). Other enriched (and depleted) TFBS can be found in Table S3.

### Evidence for DNA methylation modulating TF activity

Despite identification of interesting TFs, overall enrichment of their binding sites in promoters of DEGs was modest. The low enrichment of TFBSs in promoters of DEGs suggests that only a limited number of target genes of these TFs are hibernation-associated. As all TFBS for these genes contain at least one CpG dinucleotide, DNA methylation may modulate the impact of the TF, *i*.*e*. that a relevant fraction of putative TF target genes is not differentially expressed due to DNA methylation blocking the TF induced expression. Therefore, we set out to assess the influence of DNA methylation on gene expression, focusing on those genes with the identified motifs in their promoter regions. Methylation levels of the target genes of EGR1, MYC/MNT and CREB1 were investigated by calculating average promoter methylation as well as TFBS methylation for both DE and non-DE putative target genes. Methylation differences of DE and non-DE target genes were determined for the comparisons between hibernation stages which revealed TFBS enrichment among DE genes: *i*.*e*. (i) SE vs AE and (ii) TL vs AE for EGR1-like, (iii) SE vs AE for MYC/MNT-like and (iv) SE vs AE for CREB1. For each comparison, mixed models showed significantly higher AE promoter methylation for candidate target genes that were not differentially expressed (all P < 1.3E-06), an effect that was even more outspoken when considering methylation of the TFBS directly (Fig. 5; all P < 3.6E-05, except for MYC/MNT, P = 0.07). These results provide additional evidence for the relevance of these TFs, as the limited enrichment of their targets may be explained by the inhibitive effect of DNA methylation for non-responding TF target genes.

**Fig. 5.**
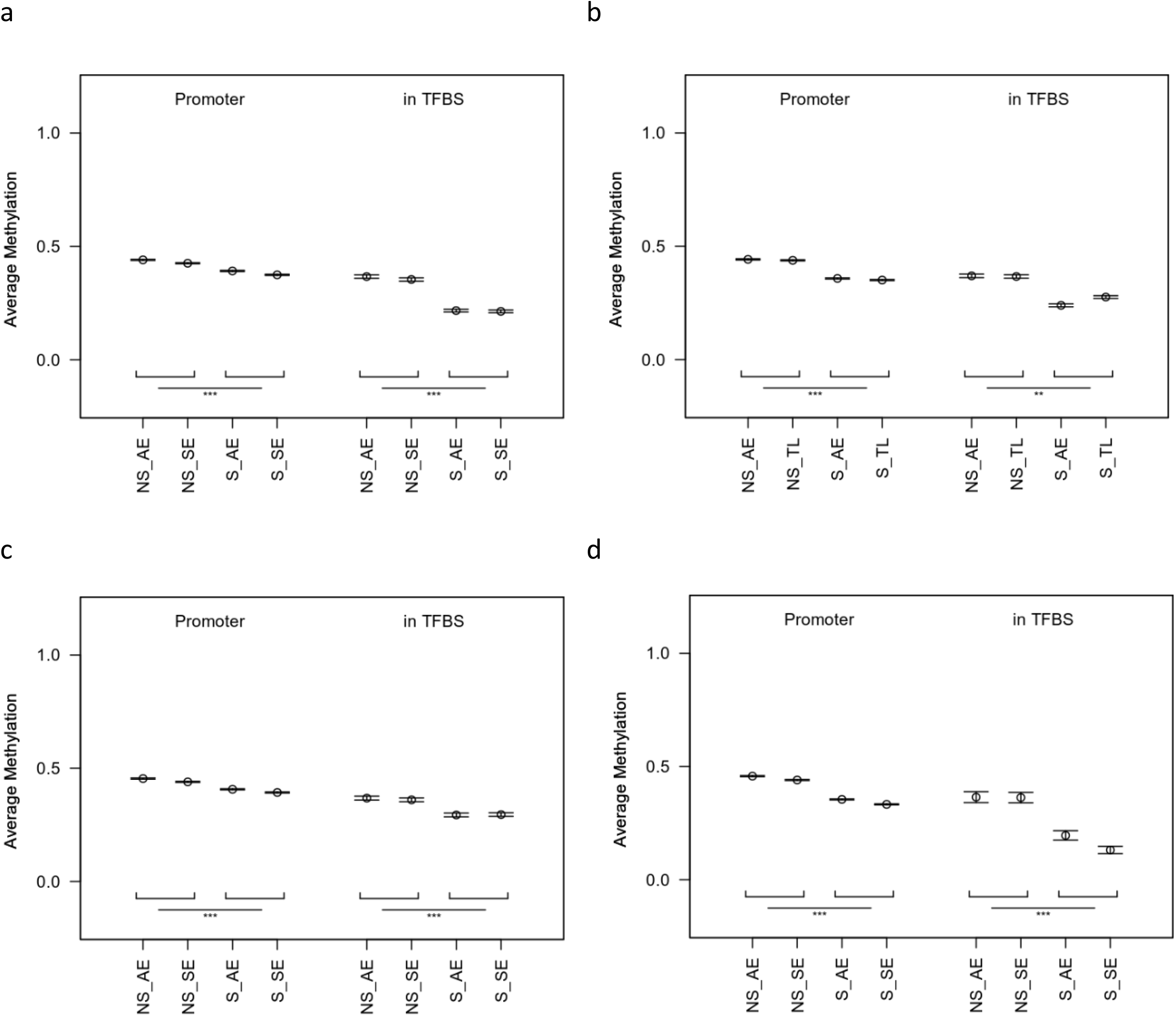
Average methylation levels of the promoter (left) and TFBS (right) for target genes identified transcription factors. NS = non-significant, S = significant. a. EGR1-like transcription factor (SE vs AE). b. EGR1-like transcription factor (TL vs AE). c. MYC/MNT-like transcription factor (SE vs AE). d. CREB1 transcription factor (SE vs AE). * * *: p value < 1*10^−3, * *: p value < 1*10^−2

## Discussion

This study is the first to investigate gene expression and its regulation in a facultative hibernator, the Syrian hamster, in an unbiased approach focusing on liver because of its critical role in metabolism, endocrinology and detoxification. Differential RNA expression analysis between all different groups revealed 619 annotated hamster genes, reflecting the dynamic nature of the transcriptome in hibernating liver. The genes group in five clusters reflecting pathways that allow the animal to accommodate long periods of physical and metabolic inactivity. The pathways regulated throughout the hibernation cycles (SE vs TL vs AE) involve metabolic processes and transcription. These pathways correspond with those previously documented by RNA-sequencing and targeted metabolomics in the liver of seasonal hibernators^6,19,24–30^ and activity assays in facultative hibernators^31^. Interestingly, our data showed three additional, prominently regulated pathways in hamster liver, i.e. the MAPK/ERK pathway, PP1 pathway and processes involved in cell ‘life-and-death’ (cell cycle, division, proliferation and death). Transcription factor binding site analysis identified 4 candidate TFs involved in regulation of gene expression of which three were differentially expressed. Moreover, methylation of promotor regions does not associate with overall differential gene expression, yet may inhibit the responsiveness of specific genes to the identified TFs.

All hibernators (partially) switch from carbohydrate to lipids as their primary fuel source during their hibernation period^4^. In keeping, our data indeed show upregulation of genes involved in gluconeogenesis and glycogenolysis during torpor. Glucose metabolism is regulated through limiting storage of glucose into glycogen by downregulation of three regulatory subunits of protein phosphatase PP1^12^, which corresponds to our data in TL and AE hamsters. The limited glucose conversion is possibly induced to maintain stable glucose levels in the hibernating hamsters. Additionally, insulin resistance during torpor leads to an impaired glucose metabolism while glycogen storage is increased^12^. Simultaneously, shifts in gene expression promote glucose retrieval from glycogen through upregulation of α-glucosidase (*GAA)*, a gene encoding an essential protein for glycogenolysis, in torpor late versus summer euthermic animals. A decrease in hepatic glycogen and upregulation of other glycogenolysis-inducing enzymes (glycogen phosphorylases), but not upregulated hepatic *GAA* expression, has been demonstrated before in hibernating animals^9^. Collectively, the regulation of genes involved in glucose-saving and glycogen-storing processes likely serve to maintain steady blood glucose throughout all hibernation stages. Moreover, the break on glycolysis observed during torpor, as demonstrated here by the strongly reduced expression of its rate limiting enzyme, *GCK1*, is released in early arousal. The reduction of glycolysis through downregulation of *GCK1* has been shown in other hibernators, such as grizzly and Asiatic black bears^5,32^.

Regulation of protein phosphorylation constitutes an important category of our identified DEGs, represented in all comparisons, including the regulation of *PP1* subunits addressed above. The downregulation of genes involved in the PP1 pathway is not unexpected, as it is known in the hibernation field that both PP1 and PP2 are regulated: decreased activity of PP1 and PP2a in torpid squirrels was reported in liver and brain respectively^33,34^, and increased PP2c activity was found in skeletal muscle, brown adipose tissue, kidney, brain and liver ^33,35^. Temperature does not seem to influence PP1 enzyme activity in hibernators, suggesting that PP1 regulation is an active process rather than passively regulated by reduced temperature^33^. We here report reduced mRNA expression of PP1 members, which could cause a decreased enzyme activity in the liver during hibernation, although studies in PP1 activity in hamster liver should be conducted to confirm this hypothesis.

Remarkably, the MAPK/ERK pathway, constituting the most prominently regulated phosphorylation cascade, is largely inhibited in both TL and AE compared to SE, mainly through upregulation of its inhibitors (multiple dual-specificity phosphatase (DUSPs) and sprouty’s (SPRYs)) in torpor. Strikingly, this strong transcriptional inhibition of the MAPK pathway has not been identified previously in hibernators. Conversely, the limited number of studies on phosphorylation status of MAPK pathway proteins show contradicting results. Increased phosphorylation of several MAPK members in liver of the obligatory hibernator Monito del Monte (*Dromiciops gliroides)* suggest activation of MAPK signaling during torpor^36^. Also in torpid Syrian hamster brains, ERK1 phosphorylation was increased. On the other hand, in the same torpid hamsters, inhibition of the MAPK pathway was observed, indicated by a strongly reduced ERK2 phosphorylation, questioning the overall activity of the MAPK kinase pathway in these animals ^37^. In torpid ground squirrel skeletal muscle, the p38MAPK activated protein kinase 2, MAPAPK2, displayed reduced activity ^38^. Our data clearly show a downregulation of the MAPK pathway in torpor and arousal at the RNA levels, which is in contrast to the torpid hamster brain, as reported previously ^37^.

Among the plethora of pathways influenced by MAPK and PP1, regulation of cell cycle arrest, cell replication and cell death constitute prominent pathways regulated at the RNA level in hibernating Syrian hamster liver. Our data provide strong evidence for the marked regulation of expression of genes involved in cell “life-and-death”, including cell division and proliferation, cell cycle arrest and apoptosis. Most studies on cell division in hibernation have been carried out on gut, showing a cessation of mitotic activity in torpor, with a progression in G_1_ phase but a block in G_2_ or S phase^39^. Also in liver of ground squirrels, cell cycle progression was suppressed during torpor as indicated by Western blot and PCR for cell cycle markers^7^. Mitotic activity and cell proliferation resume during arousals^40^. Similar to ground squirrels, our data showed an upregulation of many transcription factors and effectors involved in cell cycle regulation in the transition from TL to AE. Upregulated genes included clock gene Per1, which regulates cell growth and DNA damage control^41^, and a number of genes encoding FOS and Jun Proto-Oncogene (JUN) proteins, which are downstream components of the MAPK pathway and part of the AP-1 transcription factor process. MAPKs activate the AP-1 transcription factor complex which regulates cell growth and differentiation as a response to stress factors^42^. The inhibition of the MAPK pathway throughout torpor and partially in arousal may thus serve as a regulatory mechanism to suppress the energy costly process of cell proliferation^43^.

Our TFBS analysis identified six TFs potentially involved in upregulating gene expression in arousal. For three of these, we could demonstrate a differential upregulation at the RNA level in arousal (EGR1, MNT and MYC), whereas MYCN, MAX and CREB1 were not differently expressed. TFs are known to be regulated also at the posttranslational level and can form complexes: MYCN belongs to the same TF family as MYC and MAX is known to form heterodimers with MYC^44,45^. Interestingly, some of the identified TFs were previously found upregulated in the liver of arousing ground squirrel (MYC) and activated in muscle of the torpid brown bat and in multiple organs (including liver) of the torpid ground squirrel (CREB1)^6,23,46^, further supporting a role for these TFs in hibernation. Notably, the MAPK pathway regulates the activities of several TFs through phosphorylation, including the activation of MYC and CREB1^47,48^, revealing a possible additional role of the prominent MAPK regulation in hibernating animals. It is important to mention that the gene expression of MAPK members does not necessarily represent the activity of the MAPK enzymes. Altogether, it is indicated that the MAPK pathway plays a role during the arousal phase through upregulation of TFs and gene expression of members of the MAPK pathway.

To further explore regulatory mechanisms in hibernation, DNA methylation was measured in the liver from the hamsters. Charting of the DNA methylation on a genome-wide level did not show pronounced differences in liver between the hibernation groups, similar to ground squirrel^49^,. Contrastingly, DNA methylation in skeletal muscle of ground squirrel did show lower levels of DNA methylation in torpor late and arousal^49^. This suggests a tissue specific role for DNA methylation in hibernation. In this respect, another study showed contrasting results in livers, kidney and heart of the chipmunk by presenting hypomethylation of the USF binding site in the hibernating livers, which is responsible for upregulation of the hibernation-associated *HP-27 gene*, but showed hypermethylation in kidney and heart^50^. Tissue-specific DNA methylation could explain the differences in gene expression amongst tissues during hibernation^5,51^ Despite the similar levels of methylation genome-wide, further investigation of the methylation status showed that promoters of non-responding (not differentially expressed) target genes have significantly higher methylation. Even more strikingly, when focusing on the transcription binding site, this effect is further enhanced. We note that most candidate regulating TFs are overexpressed during arousal, possibly facilitating the switch from hibernating tissue to metabolically active tissue. Finetuning of this switch is likely subject to modulation by DNA methylation. Further characterization of these epigenetic differences could lead to interesting targets for epigenetic drugs enabling more effective organ transplantation.

Together, our study identifies that the facultatively hibernating Syrian hamster shares the regulation of key processes with seasonal hibernators, principally comprising metabolic changes, representing a preference for fatty acid oxidation and glucose-deriving processes. Our data implicate substantial expression changes in genes effectuating protein phosphorylation and discloses a profound transcriptional inhibition of the MAPK and PP1 pathways, of which the latter pathway has not been associated with hibernation before. Inhibition of these pathways seems strongly linked to cell cycle arrest and a halt in RNA transcription during torpor, both of which are restarted in early arousal accompanied with an overshoot in the expression of transcription factors. During arousal, it is suggested that restoring the expression of MAPK members leads to activation of TFs, such as MYC and CREB1, which is facilitated by hypomethylation.

Further characterization of phosphorylation cascades and their downstream factors, notably MAPK, in other hibernating species and organs is warranted to investigate the generalizability of our results.

Torpid animals tolerate hepatic ischemia following profound reductions of blood flow, whilst maintaining mitochondrial respiration, bile production, and sinusoidal lining cell viability, as well as lowering vascular resistance and Kupffer cell phagocytosis^52,53^. Our data provide further indications that administration of MAPK inhibitors might protect from cell damage by arresting cell cycle during ischemia^54^ and that MAPK regulated transcription factors may be interesting targets to avert damage by regulating cell cycle progression during organ reperfusion. Understanding molecular hibernation mechanisms may advance therapeutic approaches in medical conditions which are strongly correlated with reduced metabolism such as ischemia-reperfusion and transplantation.

## Material and methods

### Animals

Experiments were performed on male and female Syrian hamsters (*Mesocricetus auratus*) as previously described ^22^, approved by the Animal Ethical Committee of the University Medical Center Groningen (DEC 6913B) and carried out in accordance with European and Dutch legislation. Furthermore, the study is reported in accordance with ARRIVE guidelines (https://arriveguidelines.org), and all methods were performed in accordance with relevant guidelines and regulations. The following groups were included: summer euthermic (SE), torpor late (TL, torpor > 48h) and arousal early (AE, rewarming for 90 min). All hibernating animals were kept in darkness until euthanization. Arousal was induced at > 3 days of torpor by gentle handling. The animal’s activity pattern accurately identified torpor bouts of hamsters^55^, as evidenced by mouth temperature (Tm) at euthanization, being 9.0±0.9°C and 35.0±2.1°C for TL and AE, respectively, whereas SE animals had a Tm of 35.7±0.5°C. Liver was flushed with physiological salt solution, removed and snap-frozen in liquid nitrogen and stored at −80□°C.

### RNA sequencing library preparation

Total RNA was extracted from liver tissue samples of three animals per hibernation phase using Nucleospin (Machery Nagel, Düren, Germany). Concentrations were measured using Nanodrop and processed for RNA sequencing (RNAseq) by NXTGNT (www.nxtgnt.com). RNA quality was checked using a Bioanalyzer RNA 6000 nano chip assay and Ribogreen assay (Invitrogen, Carlsbad, CA, USA). 437ng RNA per sample was used for further analysis. cDNA libraries were prepared for sequencing using Truseq stranded mRNA library prep (Illumina, San Diego, CA. USA) according to protocol. Sequencing was performed on a NextSeq500 High output flow cell, generating single-end 75bp reads.

### Transcriptome sequencing and analysis

Transcriptome sequencing produced 52.6 million reads per library (range: 40.2-62.3M; Table 1). The reads were aligned using STAR aligner version 2.7.0f using genome reference MesAur1 and Ensembl 96 gene annotation. Two-pass alignment mode and gene expression quantification options were used. Out of 17,091 genes with non-zero expression, data from 11,078 genes with average expression levels above 1 fragment per million (FPM) was used for analysis. Differential expression analysis was performed using EdgeR and relied on the Benjamini-Hochberg procedure for multiple testing correction ^56^. Principal component analysis using covariance matrix was performed to visualize variation between animals (Fig. S1).

**Table 1.** Number of sequenced and uniquely mapped reads per animal. *SE= Summer euthermic, TL = Torpor Late, AE = Arousal Early*.

| **Library** | **Raw reads** | **Uniquely mapped** |
| --- | --- | --- |
| SE1 | 59,356,458 | 54,284,621 (91.5%) |
| SE2 | 55,782,947 | 51,272,445 (91.9%) |
| SE3 | 52,748,518 | 48,037,495 (91.1%) |
| TL1 | 40,161,724 | 37,095,176 (92.4%) |
| TL2 | 52,607,083 | 48,353,882 (91.9%) |
| TL3 | 53,001,874 | 48,507,248 (91.5%) |
| AE1 | 54,308,111 | 49,838,096 (91.8%) |
| AE2 | 62,274,042 | 59,230,482 (92.2%) |
| AE3 | 43,086,596 | 39,767,175 (92.3%) |

**Table 2.** Total number of differentially expressed genes (DEGs) with FDR < 0.01 between the three investigated phases of hibernation. Direction of regulation is represented against the first group in the comparison. There is a significant difference in the number of up-and downregulated genes between phases (p < 0.0001, Chi-square test). SE = Summer Euthermic, TL = Torpor Late, AE = Arousal Early.

|  | SE vs TL |  | TL vs AE |  | SE vs AE |  | SE vs TL+AE |  |
| --- | --- | --- | --- | --- | --- | --- | --- | --- |
| Total DEGs | 272 |  | 137 |  | 435 |  | 508 |  |
| Upregulated | 124 | 45% | 128 | 93% | 270 | 60% | 241 | 47% |
| Downregulated | 148 | 55% | 9 | 7% | 165 | 40% | 267 | 53% |

Expression of genes that were differentially expressed in at least one comparison (FDR<0.01) were converted into z-scores and clustering analysis was performed using pheatmap R package.

### Pathway analysis/GO enrichment analysis

Functional categories of DEGs (False Discovery Rate (FDR) < 0.01) were identified by Gene ontology (GO) analysis, categorizing DEGs in biological process (BP), molecular function (MF) and cellular component (CC) gene sets, and by Kyoto Encyclopedia of Genes and Genomes (KEGG) pathway analysis performed by g:Profiler (version e100_eg47_p14_7733820) ^57^. Identified BP pathways were visualized by means of bubbleplots using R (version 3.6.2).

### Transcription factor binding site (TFBS) enrichment analysis

Transcription factor (TF) binding site (TFBS) enrichment analysis was performed as described in Diddens at al. ^58^ for promoter regions (defined as the genomic region 2,000bp upstream to 500bp downstream of the transcription start site) of differentially expressed genes identified in transcriptome analysis (separately for up and downregulated genes per contrast). The outgroup comprised all genes that were sufficiently covered, but not significantly differentially expressed in any contrast of interest. Position weight matrices (PWM) were downloaded together with accompanying annotation from JASPAR (JASPAR 2020 server), an open source, curated database of binding motifs for transcription factors ^59^. Since transcription factors are generally well conserved and the human TFBS dataset is more complete than the murine TFBS dataset (1201 vs 529 respectively, assessed on January 15^th^ 2021), the former was selected ^60^. This has the additional advantage that any TF with enrichment in predicted binding sites for a certain contrast is also present in humans and may be an interesting target for human applications. Promoters were scanned using the PWMs via the FIMO software (v4.11.3)^61^. FIMO was run separately for each PWM due to computational limitations and all matches with P < 1.0E-4 (default setting) were retained for quantification. After quantification, enrichment was assessed using a chi-squared test per TF binding motif and Benjamini-Hochberg correction was applied to adjust for multiple testing leaving 4 candidate TFs.

### Whole genome bisulphite sequencing (WGBS)

Total DNA from liver tissue samples was extracted using Machery Nagel Nucleospin tissue kit and used for whole genome bisulfite sequencing (WGBS) by NXTGNT (www.nxtgnt.com). Three animals per hibernation phase (SE, TL and AE; the same individuals as used for RNA sequencing) were selected for sequencing. Concentration of the extracted DNA was measured using Quant-iT PicoGreen kit (Invitrogen). DNA quality was checked on a 1% agarose gel (E-gel EX Invitrogen). 500ng DNA was used from each sample for further analysis. Fragmentation was performed using Covaris model S2 to obtain fragments with a size of approximately 400 bp, followed by bisulfite conversion with the EZ DNA methylation Gold kit (Zymo Research, Irvine, CA, USA) according to manufacturer’s protocol. For library preparation, the NEBNext Ultra II DNA library prep kit (New England Biolabs, Ipswich, MA, USA) was used according to protocol, followed by sequencing on three Hiseq3000 lanes (Illumina), generating PE2×150bp reads.

### Sequence read mapping, summary and differential methylation analysis

Reads were mapped using the bowtie2 option (v 2.3.3.1) of the Bismark software (Brabraham Bioinformatics, v 0.18.1_dev) against the Syrian hamster reference genome provided by Ensembl (MesAur1.0, release 95). FastQC (Brabraham Bioinformatics, v 0.11.2) was used to assess quality of the WGBS samples, indicating good quality (both on the raw and trimmed fastq files). Trim Galore! (Brabraham Bioinformatics, v 0.1.0) was used to trim out bad quality bases (quality score < 20, default) from reads (reads trimmed shorter than 15bp were discarded). Bismark and bismark_methylation_extractor (v 0.18.0) were used to quantify CpG methylation. Finally, coverage files were compiled (bismark2bedGraph, v0.18.0) for downstream analysis.

The Bioconductor BiSeq (v 1.26.0) was used to import coverage files. Next, a gene-centered approach for WGBS data was performed, selecting all CpGs within differentially expressed genes or their promoter regions (defined as 2kb upstream and 0.5kb downstream of the transcription start site, based on Ensembl annotation). Differentially methylated regions (DMRs) were identified as described by Hebestreit et al ^62^. Initially, regions with 15 grouped CpGs (“grouped” meaning a maximum of 100bp between 2 subsequent CpGs) and at least detected in two out of 6 samples are considered. Clusters were analyzed and subsequently trimmed using default settings ^62^. Pairwise comparison between the three hibernation (SE, TL and AE) phases was performed. In order to obtain loci featuring robust methylation differences with a probable impact on expression, we only report results on DMRs displaying an average methylation difference of at least 20% and containing at least 2 CpGs (after trimming of the clusters).

### Methylation rates in TFBS and promoters with TFBS

Methylation percentages were calculated for all CpG dinucleotides that were covered at least 1 time in each sample (n = 12,405,257). For each candidate regulatory TF, CpGs in promoter regions of non-significantly DE and significantly DE target genes (i.e. genes with at least one TFBS in their promoter region) were selected. For EGR1 and MYC/MNT, closely related TFs were incorporated as well, since TFBS for these genes are nearly indistinguishable. Average methylation per promoter region or TFBS was modeled using a mixed model with the sample nested within the hibernation phase as a random effect and differential expression state (i.e. DE vs non-DE) of the gene as fixed effect. For graphical representation, the mean methylation per gene (after calculation of gene-wise average over three samples for each group) and standard error on the mean are displayed.

## Data availability

The datasets generated in this study are deposited in the GEO database under accession number GSE199817.

## Acknowledgements

This study was financially supported by a Talent PhD scholarship to MO by the Graduate School of Medical Sciences, University of Groningen, University Medical Center Groningen and two grants from the Cock-Hadders Foundation, no. 2018-30 and 2019-52.

## Author contributions

M.M.O obtained mRNA and gDNA, performed pathway analyses/ GO enrichment analyses and wrote the manuscript. L.C. performed WGBS, DMR and TFBS analysis and wrote the manuscript. V.G. performed RNA sequencing data analysis. V.A.R., J.J.B, M.G and H.R.B wrote application and performed animal experiments. T.M, M.G.R, and R.H.H supervised analyses and writing of the manuscript.

